# Automated analysis of *C. elegans* behavior by *LabGym*: an open-source, AI-powered platform

**DOI:** 10.1101/2025.08.28.672961

**Authors:** L. Amanda Xu, Hongjiang Liu, Zhaoyu Li, X.Z. Shawn Xu, Bing Ye, Yujia Hu, Elizabeth A. Ronan

**Author notes:** Correspondence (B.Y.); (E.A.R.); (Y.H.).

## Abstract

The genetic tractability, well-mapped circuitry, and diverse behavioral repertoire of the nematode *C. elegans* make it an ideal model for physiological and behavioral studies. A wide range of methods has been developed for analyzing *C. elegans* behaviors, evolving with advances in technology such as videography and computer-assisted analysis. Here, we introduce *LabGym*—an open-source, artificial intelligence (AI)-based platform we recently developed—to the *C. elegans* research community. We trained deep learning models in *LabGym* capable of automatically categorizing and quantifying multiple user-defined parameters of worm locomotion behavior in multi-worm videos with high accuracy. Furthermore, we demonstrated their efficacy in quantifying locomotion changes in aging worms. Our work offers a cost-effective, user-accessible approach to behavioral analysis in *C. elegans*.

## INTRODUCTION

Research analyzing animal behavior has provided fundamental insights into all levels of biology (Altimus et al., 2020; Pereira et al., 2020). Behavior serves as a powerful readout of an organism’s physiological state. The model organism *Caenorhabditis elegans* is an established system for behavioral genetic studies, having led to groundbreaking discoveries in neurobiology, aging, and cell biology, among others (de Bono and Maricq, 2005; Herndon et al., 2002; Xiao and Xu, 2021). Although *C. elegans* are relatively simple animals with a cylindrical body shape and a nervous system comprised of a mere 302 neurons, they are capable of exhibiting a rich repertoire of behaviors (de Bono and Maricq, 2005). Worms rely on locomotion behaviors to survive and thrive as they effectively navigate their environments, find food sources, and avoid harmful stimuli (Bargmann, 2006; Croll, 1975; Gong et al., 2019; Li et al., 2011; Pierce-Shimomura et al., 1999; Wang et al., 2016). In the laboratory setting, worm behavior is primarily two-dimensional as worms are cultivated on an agar surface, further simplifying behavioral categorization and increasing their accessibility for ease of behavioral analysis.

Methods for analysis of worm behavior continue to evolve over time with technological advances. Upon the inception of *C. elegans* as a model organism in the 1960s, initial genetic screens relied on labor-intensive manual quantification of worm behaviors (Brenner, 1974; Jorgensen and Mango, 2002). As technology developed over the decades, the implementation of videography, along with the later incorporation of computer-assisted analysis has continued to improve and standardize worm behavior analysis methods (Datta et al., 2019; Feng et al., 2006; Husson, 2012; Pokala and Flavell, 2022; Zheng et al., 2012; Hsu et al., 2009; Broekmans et al., 2016; Huang et al., 2006; Roussel et al., 2014; Wang and Wang, 2013; Javer et al., 2018; Ramot et al., 2008; Nagy et al., 2013).

Current automatic methods for analyzing *C. elegans* behavior typically combine computer vision with machine learning to estimate body posture or extract body skeletons (Alonso and Kirkegaard, 2023; Hebert et al., 2021; Liu et al., 2025; Vedantham et al., 2025; Weheliye et al., 2025). Commercial platforms, such as *WormLab* (MBF Bioscience), automate detection via computer-vision algorithms to perform centerline extraction and kinematic analysis (Roussel et al., 2014). Nevertheless, commercial software for behavioral analysis, including *WormLab*, is not open-source and comes at a financial cost, limiting user accessibility and customizability. Javer et al. (2018) established a critical open-source infrastructure for the field by introducing the OpenWorm Movement Database for large-scale data sharing and the Worm tracker Commons Object Notation (WCON), a standardized, JSON-based format for behavioral data (Javer et al., 2018). To support these resources, they developed *Tierpsy Tracker*, a software suite that bridges the gap between high-throughput multi-worm tracking and the high-resolution morphological analysis typically for single-worm studies (Javer et al., 2018). However, substantial downstream analysis is required to convert these numerical outputs into kinematic descriptors such as speed, body curvature, and turning frequency, and then into discrete behavioral states.

Recently, AI-based machine learning approaches have attracted significant attention for their potential to automate analysis of animal behavior, promising to greatly reduce users’ workload and improve reproducibility. (Goodwin et al., 2024; Goss et al., 2024; Graving et al., 2019; Hsu and Yttri, 2021; Hu et al., 2023; Kabra et al., 2013; Mathis et al., 2018; Wiltschko et al., 2015). Open source toolboxes such as *DeepLabCut* (Mathis et al., 2018) and *DeepPoseKit* (Graving et al., 2019) use Convolutional Neural Networks (CNN) to automatically track user-defined body parts for markerless pose estimation by outputting spatial coordinate values of key points. While these approaches are broadly applicable across animal species, additional tools have also been developed aimed at resolving challenges specific to behaving nematodes. For instance, *WormPose* utilizes a generative Gaussian Mixture Model (GMM) of worm postures combined with real video “textures” to create realistic synthetic training data (Hebert et al., 2021). This approach utilizes a CNN to achieve high accuracy in identifying complex, self-occluded, and coiled shapes. More recently, Alonso and Kirkegaard (2023) developed *Deeptangle*, an end-to-end deep learning approach that predicts worm centerlines directly from video clips (Alonso and Kirkegaard, 2023). This method is uniquely capable of simultaneously tracking the precise shape trajectories of thousands of swimming nematodes, even in high-density environments where frequent overlaps occur. Building on these concepts, *Deep Tangle Crawl* (DTC) was developed to further improve *C. elegans* tracking continuity and identity resolution during complex collisions and coiling events (Weheliye et al., 2025). These methods combine computer vision with machine learning to estimate body posture or extract body skeletons, and require downstream methods or tools for behavioral classification and quantification. More recently, unsupervised learning approaches have been utilized to transform raw pixel data into motion representations (Deserno and Bozek, 2025). Nevertheless, the mathematical representations obtained from these approaches still require additional steps to be translated into interpretable behavioral types, which may be difficult for researchers without extensive computational expertise.

Here, we introduce a user-friendly, AI-based approach for *C. elegans* behavioral analysis utilizing *LabGym*, a software platform that we recently developed for customizable and efficient behavioral categorization and quantification (Goss et al., 2024; Hu et al., 2023). *LabGym* leverages advanced deep learning algorithms to holistically evaluate spatiotemporal changes and overall motion patterns of target objects. We trained and validated *LabGym* models to analyze *C. elegans* behaviors, including speed, trajectory, forward locomotion, reversals, omega turns, and head-twitching, with high accuracy. Our *LabGym*-based approach is resilient to transient multi-worm collisions, maintaining individual worm tracking while also distinguishing forward from reverse movement without explicit head-tail assignment. As a proof of concept, we apply *LabGym* to automatically analyze worm behavior changes across aging. The AI-based models trained in this study are ready-to-use, and users can also train custom models to define, detect, and quantify diverse behaviors of interest with high accuracy and efficiency. All models and training datasets are shareable among users, ensuring reproducibility and transparency of behavioral analysis.

Together, this work provides the *C. elegans* research community with an open-source, user-friendly tool for automated behavioral analysis.

## RESULTS

### The pipeline for training Categorizers in *LabGym*

*LabGym* is an AI-based, open-source analysis software designed for the identification and quantification of user-defined behaviors across diverse animal species and experimental paradigms (Goss et al., 2024; Hu et al., 2023). To detect and assess user-defined behaviors, *LabGym* employs deep, customizable neural networks trained through a supervised learning approach. A detailed user manual and the source code of *LabGym* are publicly available (https://github.com/umyelab/LabGym).

To classify and quantify user-defined behaviors, *LabGym* uses the Categorizer module within its graphical user interface (GUI). This module applies a trained deep-learning model to categorize the behaviors of target animals across frame within consecutive, non-overlapping, user-specified time windows (see Methods, Goss et al., 2024; Hu et al., 2023).

To train a Categorizer, users begin by inputting training videos of animals exhibiting the behaviors of interest into the Training Module, which then extracts and identifies individual animals from the videos using one of two methods: background subtraction or the Detector submodule (Figure 1a). The Detector submodule was introduced in the recent major release of *LabGym* (*LabGym2*) (Goss et al., 2024). Adapted from *Detectron2* (Wu et al., 2019), a deep-learning instance-segmentation package, the Detector is an object-tracking component that utilizes a mask region-based convolutional neural network to generate segmentation masks for user-defined objects. This approaches enables robust and efficient object detection even in the presence of background noise and reduces identity switching when animals come into close proximity (Goss et al., 2024).

**Figure 1.**
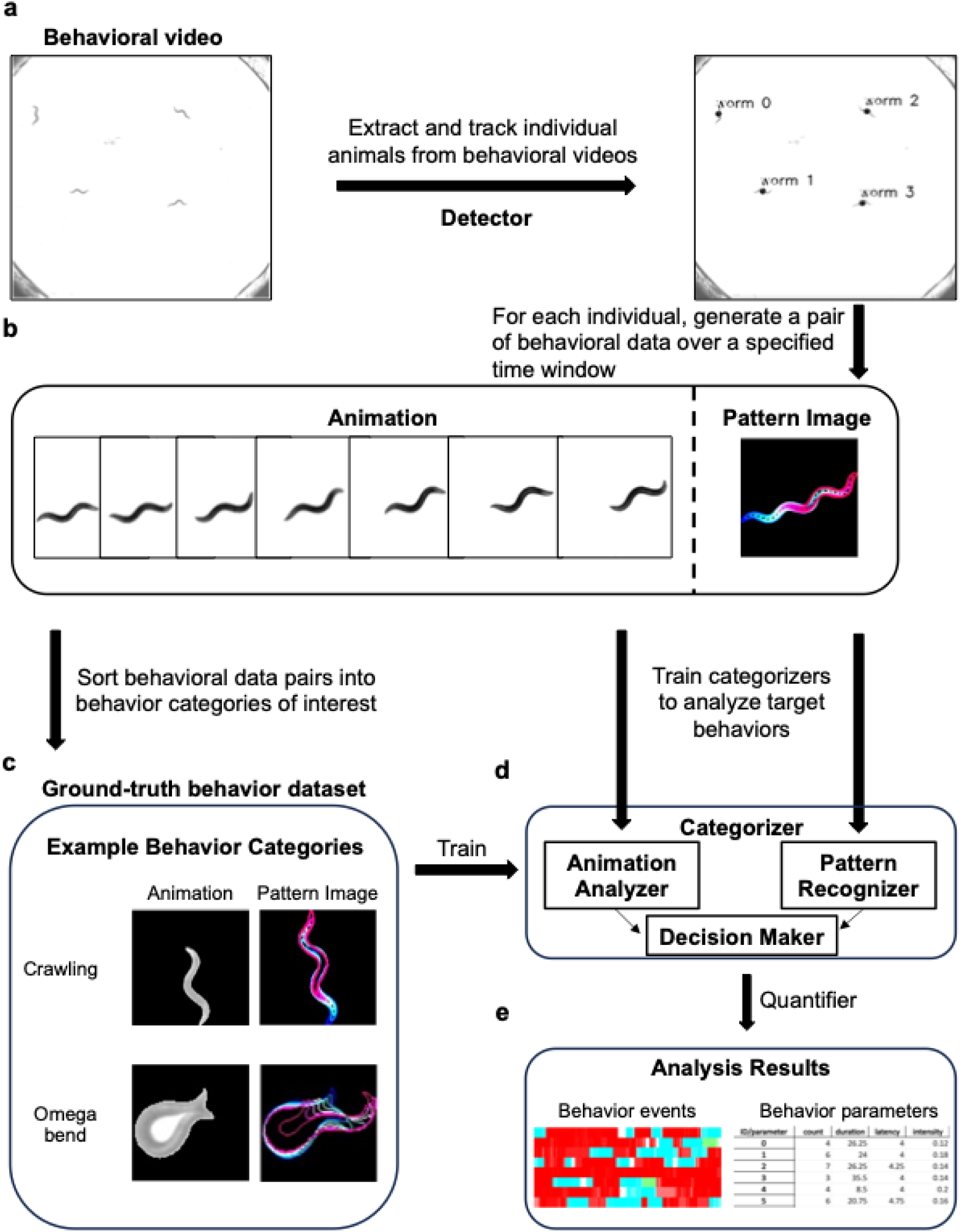
The pipeline for training deep learning models in *LabGym* for *C. elegans* behavioral analysis. **(a)** Tracked individual worms are extracted from each frame of a behavioral video using the Detector module. **(b)** At each frame of a behavioral video, *LabGym* outputs a pair of behavioral data for each worm over a specified time window: an Animation of the behavioral movement and its corresponding Pattern Image. **(c)** To train the Categorizer, specify behaviors of interest by sorting the generated pairs of behavioral data into user-defined behavioral categories. The Categorizer will utilize both the animation and pattern image to identify and quantify the target behaviors. **(d)** A trained Categorizer analyzes both the Animation (via the “Animation Analyzer”) and the Pattern Image (via the “Pattern Recognizer”) of tracked worms to categorize their individual behaviors over a specified time window. **(e)** After behavioral categorization by the Categorizer, the Quantifier evaluates categorization data and animal position to compute quantitative metrics of behaviors exhibited by each worm over any user-specified duration of an analyzed behavioral video.

Once individual animals are identified, *LabGym* generates a pair of behavioral data for each individual across consecutive, user-defined time windows. Each data pair consists of a pixelated “animation” showing the animal’s appearance and postural changes as they appear in the input video, along with a corresponding “pattern image” that encodes the animal’s movement pattern from the beginning to the end of the movement using a color gradient from blue to white and then pink (Figure 1b). This dual representation captures both broader locomotor patterns and fine-scale postural changes.

Following the generation of behavior data pairs, users sort the data pairs into user-defined behavior categories (by separating them into individual folders named by behavior type) to create the ground-truth dataset for training the Categorizer (Figure 1c). As *LabGym* employs a supervised learning approach, this annotation step is essential for providing the ground-truth examples that enable the Categorizer to learn to recognize each target behavior type.

After the ground-truth behavior dataset is established, it is used to train a Categorizer, which employs deep neural networks (the Animation Analyzer and the Pattern Recognizer) to process the two types of behavioral information extracted from each frame of a video (Figure 1d; see Methods). The Animation Analyzer focuses on the time-varying data in the “animation”, extracting details about the animal’s posture and body shape at each frame. In contrast, the Pattern Recognizer examines a condensed 2-D “pattern image” that captures the overall motion path of the animal across consecutive frames, allowing the Categorizer model to interpret movement changes over time. These two analytical streams are then integrated by the Decision Maker, which assigns a behavior category to each video frame and calculates the event probability—a percentage representing the likelihood that the given behavior occurred at that frame (Figure 1d).

Following behavior categorization, the Quantifier module integrates behavior category labels from the Categorizer and positional information of each tracked individual to calculate detailed quantitative measurements of behaviors exhibited at each frame, which are outputted into Excel files organized by individual animal and behavior type (Figure 1e). Additional outputs include raster plots color-coded by behavior category and corresponding probability, to enable temporal visualization of all behaviors exhibited by each individual over the analysis window (Figure 1e). *LabGym* also generates annotated videos of the original behavioral recordings, where each individual is labeled frame-by-frame with its classified behavior and corresponding event probability (Videos S2 and S3).

### *LabGym* accurately analyzes user-defined *C. elegans* locomotion behavior

To test if *LabGym* can be applied to analyze *C. elegans* behavior, we attempted to train Detectors and Categorizers for identifying and quantifying worm locomotion, a prominent *C. elegans* behavior. Worms spend the majority of their time crawling forward, with occasional reversals typically followed by omega bends to reorient their movement (Parida, 2022; Croll, 1975; Pierce-Shimomura et al., 1999). However, unlike other more complex organisms, for example, insects and vertebrates, worms are relatively uniform in morphology (Altun et al., 2009). Such morphological simplicity poses challenges for automated analysis of *C. elegans* behaviors, such as reliably tracking individual worms during self-overlap (e.g., omega bends) or contact with other worms (e.g., multi-worm collisions), as well as differentiating the head from the tail of worms. Therefore, besides basic parameters (e.g., trajectory and speed), we primarily focused on analyzing parameters such as forward and reverse crawling, which typically require accurate head/tail distinction, and omega bends. We also attempted to analyze immobility and head-twitching, two behavioral states infrequent in younger adults but increasingly common in aged worms (see below).

In this study, we focused on tracking individual-worm behavior profiles within multi-worm assays. To do so, we trained a Detector named *worm detector 9*, using the Train Detectors submodule embedded within the *LabGym* Training Module. Individual worms from multi-worm images were annotated and augmented in Roboflow. We then trained a Categorizer, *worm locomotion categorizer 22*, to identify behavioral types of N2 worm locomotion (Figure 2a; Video S1; Table S1), including forward crawling, reverse crawling, omega bend, immobile, and twitching (Figure 2a). This categorizer was trained with behavioral data extracted from videos of locomoting, 1-day old N2 worms, and locomoting N2 worms across age, and is thus potentially generalizable to various experimental conditions. We achieved an overall Categorizer accuracy of 0.90, as estimated by the F1 score (see Equation 2 in Methods and Hu et al., 2023). The confusion matrix used to calculate the precision, recall, and F1 score shows strong agreement between the actual and predicted behavioral types (Figure 2b). It should be noted that the five selected categories represent only a subset of locomotion behavior states that *LabGym* can analyze, as its Categorizers are fully customizable and allow experimenters to define behavior categories according to their research needs.

**Figure 2.**
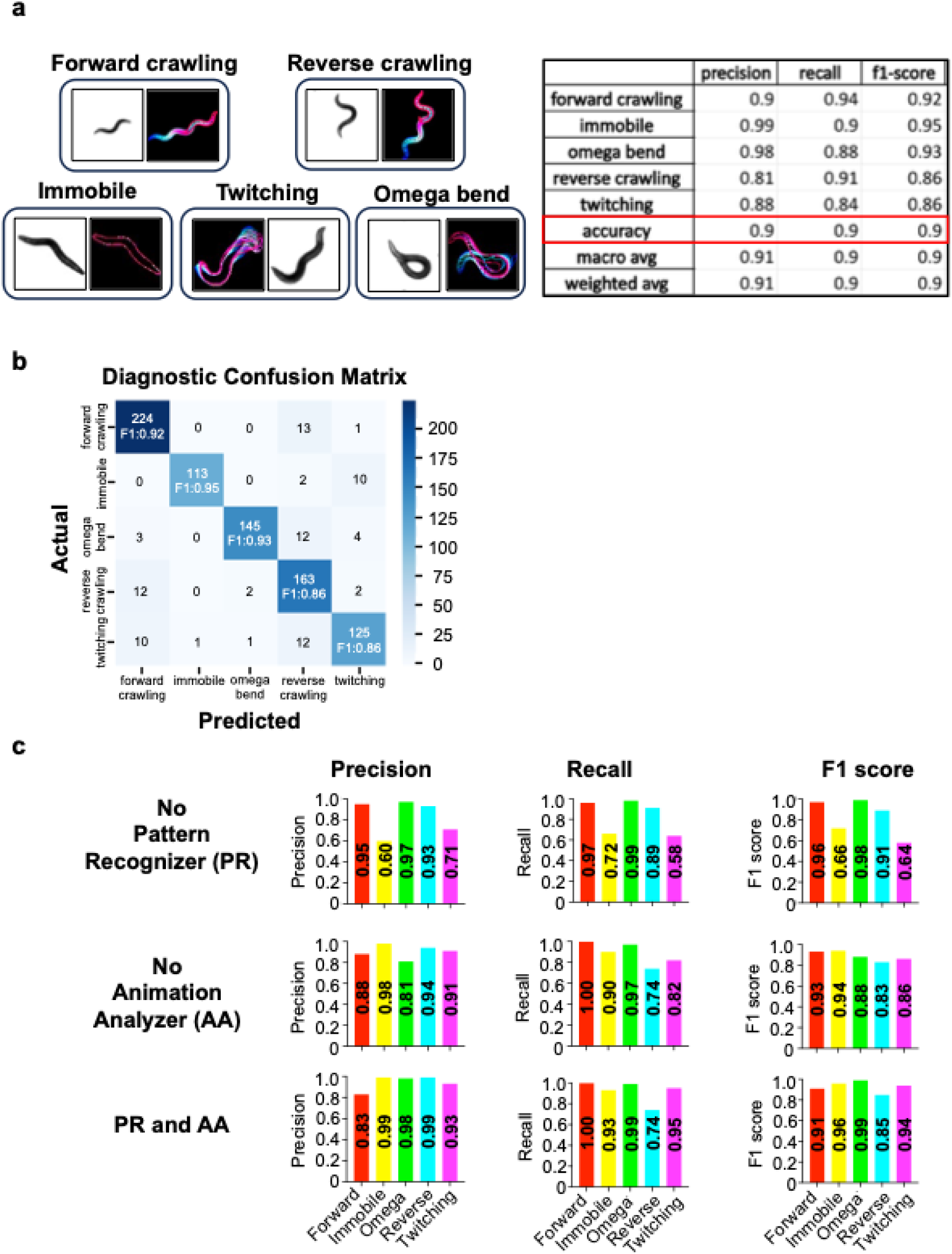
Incorporation of both the Pattern Recognizer and Animation Analyzer optimizes the categorization accuracy of *C. elegans* behaviors by *LabGym*. **(a)** Representative behavior example pairs used to train the Categorizer *worm locomotion categorizer 22*. The categorization precision, recall, and f1-score of each target behavior, as well as the overall categorization accuracy of each metric, are summarized in the table. The Categorizer training dataset included behavior examples of worms of different age, with breakdown per age as follows: 1-day-old (1,478 behavior examples), 3-day-old (115 behavior examples), 5-day-old (141 behavior examples), 7-day-old (249 behavior examples), 9-day-old (230 behavior examples), 11-day-old (169 behavior examples), 13-day-old (242 behavior examples), and 15-day-old (262 behavior examples). **(b)** Confusion matrix detailing the categorization accuracy of the Categorizer. Elements in the diagonal represent correct predictions, whereas elements in the off-diagonal represent incorrect predictions where the model confuses its behavioral categorization. The confusion matrix was generated from the same testing dataset used to generate the summary table (Figure 2a), with f1-scores in the diagonal of the matrix aligning with those in the table. **(c)** Ablation analyses demonstrate that the incorporation of both the Pattern Recognizer and Animation Analyzer modules of *LabGym* optimizes the categorization accuracy of the Categorizer. Bar graphs depict the precision, recall, and f1-score of analysis accuracy for three versions of the Categorizer *worm locomotion categorizer 22*: one version with the Pattern Recognizer ablated (top), one version with the Animation Analyzer ablated (middle), and one version with the inclusion of both modules (bottom).

We also tested the importance of Pattern Recognizer (PR) and Animation Analyzer (AA) in the *LabGym* architecture by conducting an ablation analysis (Figure 2c). Consistent with our previous report of *LabGym*, exclusion of AA leads to lower precision, recall, and F1 scores across behavioral types than inclusion of both AA and PR, whereas exclusion of PR results in the lowest precision, recall, and F1 scores overall (Figure 2c). These results demonstrate *LabGym*’s holistic assessment approach, as the involvement of both the PR and AA contributes to behavioral analysis accuracy by *LabGym*. Exclusion of PR led to lower F1 than exclusion of the AA, suggesting that the PR is more efficient than AA to capture the worm behaviors in our dataset, which is also consistent with the conclusion in our previous report (Hu et al., 2023).

The assessments above reflect *LabGym*’s performance on a ground-truth testing dataset consisting of sorted example pairs that were not used to train the Categorizer (see Methods and Hu et al., 2023). To directly assess the practical accuracy of the Categorizer in identifying and quantifying *C. elegans* behaviors, we compared the *LabGym*-computed behavioral outputs with human (i.e. manual) annotation of N2 worm locomotion (Figures 3a-d). Side-by-side raster plots of *LabGym* and manual analyses revealed strong agreement between the two methods (Figure 3a). We then calculated the percentage of frames in which the behavioral events computed from *LabGym* and manual analyses matched for each worm, and averaged across individuals, yielding an overall accuracy of 90% (Figure 3b; see Methods). This indicates that *LabGym* is estimated to successfully analyze 90% of behavioral events identified by human vision.

**Figure 3.**
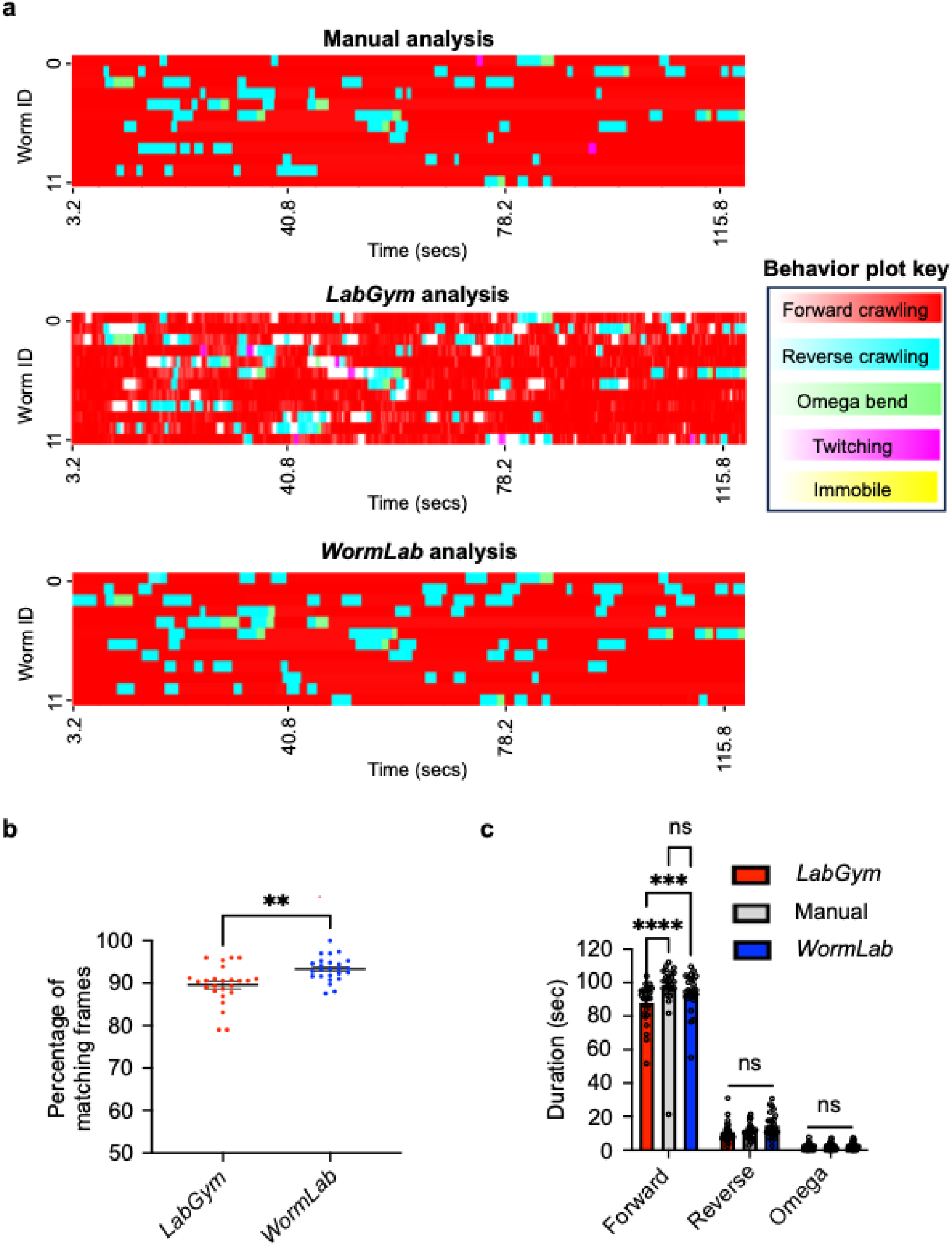
*LabGym* analyzes *C. elegans* behaviors with accuracy. **(a)** Raster plots comparing the categorization of behaviors of a representative subset of wildtype N2 worms (n = 12) by human annotation (top), *LabGym* (middle), and *WormLab* (bottom) per frame (every 0.25 seconds) over a 2-minute time window. X-axis: time (in seconds); Y-axis: each row corresponds to an individual worm. Behavior plot key: color bars indicate behavior categories and their corresponding colors in the raster plot. White spaces indicate uncategorized behavioral events. The length of a colored behavioral event in the raster plot correlates with event duration (in seconds), and the color intensity of a behavioral event in the *LabGym* raster plot correlates with its *LabGym*-computed probability (from 0 to 1.0). **(b)** Quantitative benchmarking of the analysis accuracy of *LabGym* to *WormLab* in an enhanced sample of wildtype N2 worms from 4 different plates and days. Paired t-test. Side-by-side scatter dot plot, n = 25 per category. **p<0.01 (Left): The true accuracy of the *LabGym* Categorizer, determined by the percentage of matching behavioral events between *LabGym* and manual analysis for each individual worm in the raster plot. Scatter dot plot, n = 25. Mean ± SEM (0.90 ± 0.009). (Right): The true accuracy of *WormLab* analysis, determined by the percentage of matching behavioral events between *WormLab* and manual analysis for each individual worm. Scatter dot plot, n = 25. Mean ± SEM (0.93 ± 0.006). **(c)** Event duration analysis by the *LabGym* Categorizer benchmarked to *WormLab*. Behavior event durations calculated from *LabGym* and *WormLab* were individually compared with manual analysis. Two-way ANOVA with Bonferroni. Grouped bar graph, n = 25 per behavior category. ****p<0.0001, ***p<0.001, and ns: not significant.

To compare the analysis accuracy of *LabGym* with commercial standards, we analyzed the same dataset using *WormLab*, a computer vision skeleton-based software widely used in the community for worm behavioral analysis (Figures 3a-d). The percentage of matching behavior event frames between manual and *WormLab* analysis outputs was computed to be 93.3% (Figure 3b), comparable with *LabGym*’s accuracy of 90%. This suggests that these two tools perform similarly in categorizing worm behaviors as identified by human vision. Although the accuracy percentage of *LabGym* is slightly lower than that of *WormLab* (Figure 3b), increasing the size of the training dataset is expected to further increase *LabGym*’s accuracy. It is important to note that *WormLab* requires manual intervention to verify head/tail assignment before accurate behavioral quantification can be achieved. In our tests, *WormLab*’s internal head/tail assessment feature spontaneously misassigned an average of 5.2 times across videos (see Methods), such that its calculated accuracy of 93.3% was only attained after manual correction of these errors. In contrast, *LabGym* bypasses the need for head/tail identification, automatically quantifying behaviors without user intervention.

To further evaluate the Categorizer’s performance, we compared the total durations (in seconds) of each behavioral parameter computed by manual scoring and *WormLab* with *LabGym*. The *LabGym*-calculated duration of reverse crawling and omega bending behavioral types did not significantly differ from manual analysis or *WormLab* (Figure 3c).

We did observe a slight decrease in *LabGym*-computed cumulative duration of forward crawling compared to both *WormLab* and manual scoring (Figure 3c). This underestimation likely stems from the specific uncertainty thresholds and minimum behavior length episodes selected during the Categorizer training, which were tuned to prioritize high precision in identifying discrete behavioral events (i.e. reversals and omega bending) at the expense of sensitivity of all forward-crawling bouts (see Discussion). Despite this conservative output of forward locomotion, the close alignment of *LabGym*, *WormLab*, and manual analysis across the remaining behavior categories provides further validation of the Categorizer’s accuracy.

We further performed a manual audit during multi-worm collisions for individual worms to determine *LabGym’*s accuracy at maintaining identity retention and behavioral classification during these events. Though our *LabGym* model achieved an initial ∼70% accuracy rate during multi-worm collision events (Video S3), those few errors that did arise from behavioral misclassification or loss of animal detection during multi-worm contact were automatically resolved by *LabGym* relatively quickly in post-collision frames. Moreover, as Categorizers analyze behaviors in a frame-by-frame manner and miscategorizations that may occur in prior collision frames do not affect categorizations in subsequent frames post-collision, *LabGym* is capable of self-resolving collision-caused detection error. Further improvements to the classification accuracy during multi-worm collisions can likely be achieved by retraining the Detector and Categorizer on an expanded dataset that includes more image/behavioral examples containing multi-worm collisions. This highlights a key strength of *LabGym*: its categorization accuracy is achieved in a fully automated manner, thereby reducing user bias, saving analysis time, and improving reproducibility. In summary, we show that *LabGym* is capable of accurately analyzing user-defined *C. elegans* locomotion behavior. To aid users in step-by-step use of our *LabGym* pipeline to analyze *C. elegans* behaviors, we have included a “Quick-Start” tutorial (Supplemental File 1). Extensive additional resources to facilitate implementation of *LabGym* are also available on LabGym.org, including an extended user guide, video tutorials, an AI-troubleshooting companion, and a community Discord server to provide real-time support.

### *LabGym* captures behavioral changes in aging *C. elegans*

Having demonstrated the efficacy of *LabGym* in analyzing *C. elegans* locomotion behavior with high accuracy, we next sought to apply this approach to analyze biologically meaningful behavioral paradigms. One such application is in the context of aging, for which *C. elegans* is a well-established model system. The worm’s short lifespan of approximately three weeks, genetic tractability, and well-characterized developmental timeline provide a powerful platform for investigating age-dependent changes in locomotion behavior (Liu et al., 2013; Son et al., 2019).

Worm locomotion has been shown to decline rapidly with age. The forward crawling speed in exploratory behaviors typically peaks in early adulthood (Day 1-5) and progressively declines over time (Liu et al., 2013). In late age (Day 11 and beyond), functional declines in the motor nervous system and muscle lead to marked reductions in overall locomotion activity, increased immobility (Liu et al., 2013; Herndon et al., 2002; Herndon et al., 2018). Aging studies are longitudinal and require analyses at multiple time points across lifespan, making behavioral analysis labor-intensive. Therefore, automated, AI-assisted detection of age-related behavioral changes would be a valuable tool in *C. elegans* aging studies. We thus applied our *LabGym* approach to analyze locomotion behavior in aging worms.

Using Categorizer *worm locomotion categorizer 22*, we analyzed behavioral videos of aging worms across adult lifespan, repeating every other day until Day 15 adulthood. Older worms were not analyzed, as they began to die and exhibited little to no locomotion activity. For each video, *LabGym* automatically generated trajectory plots color-coded by individual worm, which revealed a decline in exploratory locomotion across age (Figure 4a), consistent with previous findings (Liu et al., 2013). Similar age-related reductions were also apparent in trajectory plots colored by time progression (Figure S1). These consistencies suggest that our Categorizers reliably capture age-associated changes in worm locomotion.

**Figure 4.**
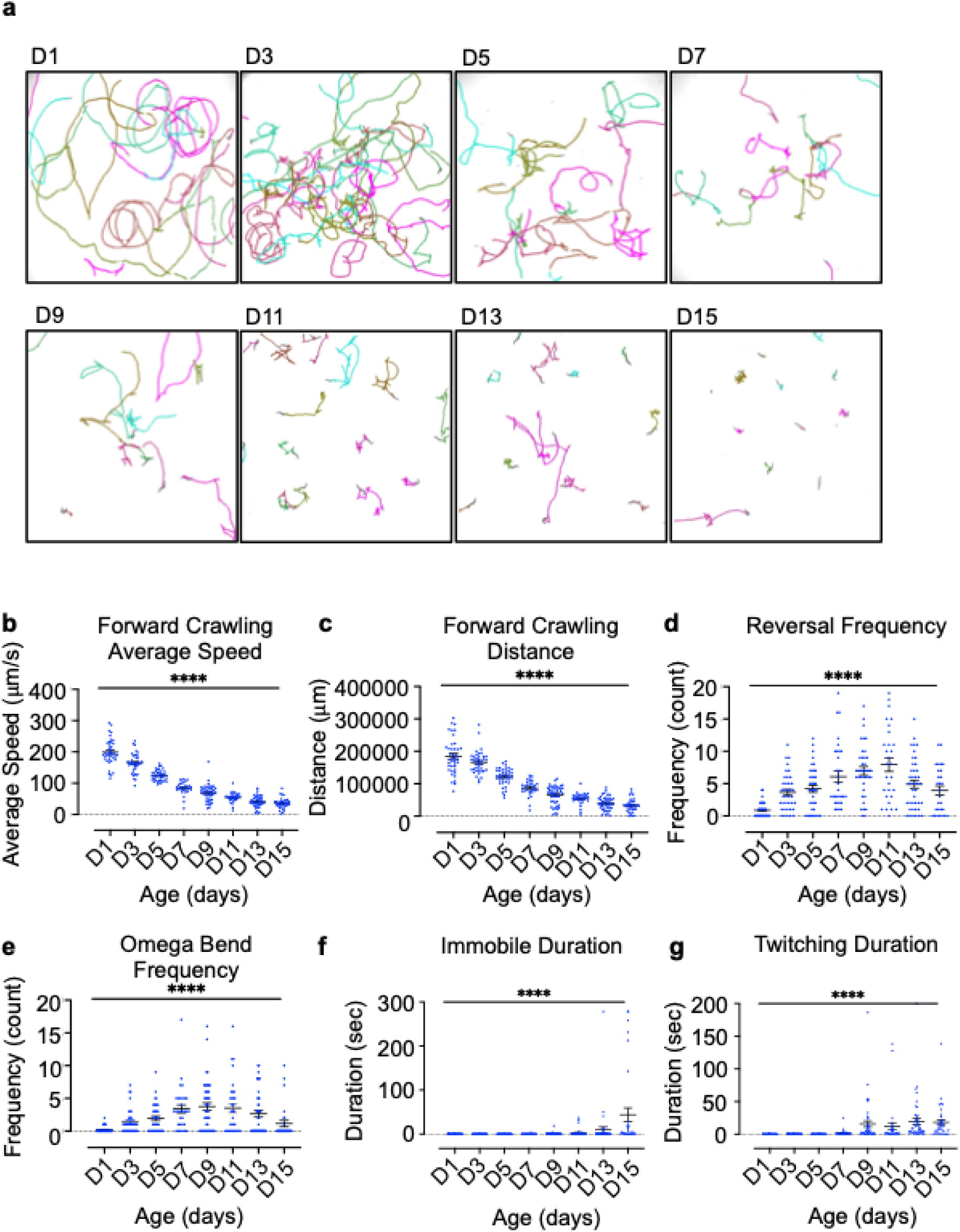
*LabGym* captures locomotion changes in aging worms. **(a)** Trajectory of a representative subset of tracked N2 worms from Day 1 (D1) to Day 15 (D15) adulthood. Each colored trace represents an individual worm. Worms were tracked over a 4.5-minute interval in each image. **(b) - (g)** Parameters of behavioral changes exhibited by aging worms from Day 1 (D1) to Day 15 (D15) adulthood, as calculated by *LabGym*. Samples from each day age was extracted from 3 behavioral videos. One-Way ANOVA. Scatter dot plot, D1 (n = 29), D3 (n = 39), D5 (n = 39), D7 (n = 30), D9 (n = 46), D11 (n = 31), D13 (n = 43), and D15 (n = 34). ****p<0.0001.

To further evaluate *LabGym*’s accuracy in assessing age-related behavioral changes, we quantified changes in individual behavior metrics across age (Figures 4b–g) and evaluated whether observed trends are consistent with prior research. Forward crawling speed (Figure 4b) and distance (Figure 4c) declined with age, in congruence with observed decreases in worm exploratory locomotion over time (Glenn et al., 2004; Liu et al., 2013). Reversal and omega bends frequencies peaked in mid-adulthood (Figures 4d and 4e), in line with reports of increased reorientation during these ages (Glenn et al., 2004; Wirak et al., 2022). In late-age adults, movement became more erratic and less coordinated, marked by elevated levels of immobility (Figure 4f) and bouts of head-twitching (Figure 4g). This may reflect a physiological shift from locomotion to near-complete immobility at late life, likely driven by motor nervous system decline and muscle degeneration (Glenn et al., 2004; Liu et al., 2013; Guo et al., 2012). In summary, we developed a Categorizer capable of detecting age-dependent behavioral changes in concordance with prior literature. This demonstrates the utility of *LabGym* as a robust tool for automated behavioral analysis in *C. elegans* aging studies.

## DISCUSSION

In this study, we demonstrate *LabGym* as a tool to automatically analyze *C. elegans* behavior. We trained *LabGym* models for analyzing worm behaviors across adult lifespan, and demonstrated its feasibility in detecting behavioral changes across aging. This work represents the first application of the *LabGym* framework to *C. elegans* research, providing a new and open-source approach for automated quantification of worm behavioral states using a holistic, skeleton-free methodology. Our pretrained models and the datasets for training them are publicly available and ready to use, allowing researchers to use them directly for deep phenotyping worm behavior and for benchmarking their own tools. Beyond the behavior categories and experiments described here, the accessibility and customizability of *LabGym* enable users to feed their own data for training it to be used for analyzing their data. This enables users to train these or their own models for analyzing *C. elegans* behaviors of interest across diverse backgrounds and settings, opening new avenues in biological research.

Distinguishing forward from reverse locomotion poses a particular challenge in *C. elegans* behavioral classification due to the morphological similarity between the worm’s head and tail, especially under low-magnification recording. *LabGym* does not require users to manually designate the head versus tail prior to behavioral classification. To reliably resolve forward versus reverse locomotion, we optimized the *LabGym* Categorizer parameters by implementing a 5% uncertainty threshold and a minimum episode length of 3 frames. These settings were selected via an empirical optimization process that maximized the accuracy of our Categorizer in detecting discrete locomotion behaviors, while prioritizing precision. As identification of reverse crawling carried more weight than forward crawling in our analysis, we adjusted the Categorizer to be more precise in identifying reverse crawling at the cost of losing some sensitivity in identifying forward crawling. As a result, our *LabGym* tuning parameters resulted in a slight underestimation of total forward-crawling duration (Figure 3c) compared to manual analysis and *Wormlab*. Our *LabGym* model’s underestimation is likely due to the filtering of forward crawling events that were either ambiguous (fell out of the uncertainty criteria) or too short (fell out of the minimum length criteria of <3 frames). Users interested in further improving *LabGym*’s accuracy at detecting forward locomotion events could modify the minimum length criteria of forward versus reverse crawling.

Additional challenges in behavioral classification can arise when worms adopt overlapping body postures (e.g., omega bends) and conformations (e.g., multi-worm entanglement). Although high-magnification videography might improve the resolution of worm morphology, it requires more expensive equipment and typically accommodates only a small number of worms, thereby limiting feasibility and experimental throughput. Notably, *LabGym* can precisely quantify behaviors even under low-magnification and during multi-worm assays.

The *LabGym* analysis accuracy of identity retention and behavioral classification of individual worms during multi-worm collisions can be further improved with an enhanced Detector training dataset that includes more examples of colliding worms. While the dataset used to train *worm detector 9* was annotated and augmented in *Roboflow* (Dwyer, B. et al., 2026), the recently developed annotation tool EZannot (https://github.com/yujiahu415/EZannot) provides a more streamlined, *LabGym*-embedded platform and automatically generates up to 135x augmented versions of each training image. Future implementation of EZannot in Detector training may improve *LabGym*-based detection accuracy. This may be especially beneficial for population assays or studies focused on multi-worm interactions.

Recent research has moved toward integrating tracking with advanced behavioral analysis. Liu et al. (2025) introduced a high-speed framework combining YOLOv8 with the ByteTrack algorithm, enhanced by a Convolutional Block Attention Module (CBAM) (Liu et al., 2025). This platform enables real-time quantification of parameters such as bending angles, roll events, and omega turns. In this framework, behaviors are identified by computational approaches, such as calculating curvature integrals along the movement trajectory. Zhan et al. (2026) developed a specialized pipeline using image processing techniques—including threshold segmentation and thinning—to skeletonize nematodes and extract their movement centerlines (Zhan et al., 2026). By employing a CNN network to localize the head and tail, the system uses mathematical approaches to classify and quantify head swings and omega turns. Both Liu et al. and Zhan et al. are methodologically different from *LabGym*, which identifies behaviors through supervised learning and does not rely on pose estimation or the extraction of specific mathematical parameters to define behavioral types. In contrast, the newer frameworks described by Liu et al. and Zhan et al. use extracted parameters (like curvature and segment angles) as the primary basis for behavioral identification (Liu et al., 2025; Zhan et al., 2026). Therefore, *LabGym* complements this landscape by applying a holistic assessment approach to evaluate an animal’s overall motion patterns and spatiotemporal changes (Hu et al., 2023; Goss et al., 2024). It bypasses the need for explicit head/tail assignment, can quantify omega bends, can accurately resolve individual worms during multi-worm collisions, and can automatically correct the errors if worms are misassigned following collisions. We attribute this success to two main factors. First, *LabGym*’s holistic assessment approach evaluates an animal’s overall motion patterns and spatiotemporal changes (Hu et al., 2023; Goss et al., 2024). Second, the deep convolutional neural networks underlying *LabGym* are trained via a supervised learning approach (Hu et al., 2023; Goss et al., 2024). The human-integrated component of *LabGym* capitalizes on the strengths of human vision and cognition to ensure that training datasets are validated by experimenters themselves.

A limitation of this supervised approach is the manual effort needed during model training, demanding greater initial user investment compared to unsupervised methods. Nevertheless, once Categorizers are trained, they can be readily applied across experiments, making the initial investment worthwhile. To further reduce user labor and improve utility, *LabGym* incorporates user interfaces that streamline the training process, such as modules for sorting behavioral examples during Categorizer training.

The Analysis module of *LabGym* currently outputs 13 quantitative measurements after behaviors are classified. While we only focused on a subset of locomotion parameters in this study to demonstrate the utility of *LabGym*, users can certainly analyze additional measurements and even define new measurements. In addition, *LabGym* has the potential to be applied to more complex locomotion-based behaviors, such as sensory behaviors (e.g., chemotaxis, phototaxis, phonotaxis, and electrotaxis) (Bargmann, 2006; Iliff et al., 2021; Liu et al., 2010; Wang et al., 2023; Ward et al., 2008; Gabel et al., 2007), as well as interactive behaviors like social feeding and mating in multi-worm contexts (de Bono and Bargmann, 1998; Liu and Sternberg, 1995). In summary, our work highlights not only the feasibility, but also the potential, of *LabGym* for behavior-based applications in *C. elegans* research. The open-source availability and user-friendly nature of the platform lower the technical barrier of its use, which may encourage researchers to customize and apply *LabGym* to their own behavioral studies.

## MATERIALS AND METHODS

### *C. elegans* strains cultivation and maintenance

The wildtype *C. elegans* strain N2 was used for all experiments. Worms were maintained at 20°C on nematode growth medium (NGM) with *Escherichia coli* strain OP50. Age-synchronization was performed by transferring L4-stage worms onto nematode growth medium (NGM) plates seeded with OP50 and allowed to develop to Day 1 of adulthood (D1). All experiments were performed on adult worms.

### *C. elegans* behavioral assays and videography

To assess behavior in D1 young adult worms, each trial consisted of transferring 5-10 age-synchronized D1 worms to an NGM testing plate without OP50, using an eyelash to minimize stimulation. Worms were allowed to acclimate for 2 minutes prior to behavior recording. Behavioral videos were recorded using the *WormLab* tracking system (MBF Bioscience) with an AVT Stingray F504B camera equipped with an AF Micro-Nikkor 60 mm lens (C-mount) mounted above the assay plates. Worms were illuminated via transmitted LED illumination from an MSCOP-004 LED illuminator (BMF Bioscience) positioned beneath the plate. For D1 adult behavior, videos were recorded at 2056 x 2056 pixels and 7.96 frames/s, corresponding to a field of view of approximately 20 × 20 mm.

For experiments examining age-related behavior changes, ∼300 age-synchronized L4-stage worms were transferred onto NGM maintenance plates containing 50 µM fluoro-2′-deoxyuridine (FuDR) to prevent progeny production. Behavior was recorded using the *Wormlab* tracking system described above on days 1, 5, 7, 9, 11, 13, and 15 of adulthood. At each time point, 10–25 worms were placed on unseeded NGM testing plates and behavior was recorded at 2456 x 2052 pixels and 7.5 frames/s, corresponding to a field of view of approximately 25 x 21 mm. Worms were not reused after behavioral recordings.

### Computational hardware

The computational procedures in this study were performed on the following computer system: Dell Enterprise Server with Microsoft Windows 11 Education, 3.10 GHz Intel Xeon w3-2435 with 64 GB RAM, NVIDIA RTX 4500 Ada Generation with 24 GB VRAM.

### Behavioral video processing

Behavioral videos (video format: .avi) to assess D1 adult behaviors were downsized to 3 frames per second using the *LabGym* Preprocess Videos submodule in the Preprocessing Module. During analysis, the framesize was further downsized using the Analysis Module of *LabGym*.

For experiments examining age-related behavior changes, videos (video format: .avi) were downsized to 2056 x 1716 pixels and 4 frames per second using the *LabGym* Preprocess Videos submodule in the Preprocessing Module. During analysis, the framesize was further downsized using the Analysis Module of *LabGym*.

### The architectures for neural networks

#### Detector

The architecture of the Detector is based on Mask R-CNN (Mask Regional Convolutional Neural Network) with a ResNet-50 backbone.

#### Animation Analyzer

The Animation Analyzer consists of convolutional blocks wrapped with time-distributed layers and followed by recurrent layers implemented using long short-term memory (LSTM). Two architectures are implemented: VGG-like and ResNet-like.

Seven levels of model complexity are provided so that users can choose an architecture appropriate for their dataset. Levels 1-4 follow VGG-like architectures containing 2, 5, 9, and 13 convolutional layers (Conv2D), respectively. Each convolutional layer is followed by a batch normalization (BN) layer. Max pooling (MaxPooling2D) layers are inserted after the second BN layer in level 1; after the second and fifth BN layers in level 2; after the second, fifth, and ninth BN layers in level 3; and after the second, fifth, ninth, and thirteenth BN layers in level 4. Levels 5-7 follow ResNet-like architectures, corresponding to ResNet-18, ResNet-34, and ResNet-50, respectively.

After the convolutional operations, the outputs at each time step are flattened into 1-D vectors and then passed to the LSTM. The outputs of the LSTM are subsequently fed into the Decision Maker submodule.

Together, these options provide a range of network architectures, from a simple 2-layer VGG-like model to a more complex ResNet-50-like architecture, allowing users to select an appropriate model complexity for their datasets.

#### Pattern Recognizer

The Pattern Recognizer consists of convolutional blocks that take 3-D tensors (width × height × color channels) as inputs. The width and height are user-definable, while the number of color channels is fixed at three (RGB: red, green, and blue) because the colors in pattern images encode the temporal sequences of animal movements.

Seven complexity levels are also available for the Pattern Recognizer to accommodate datasets of varying sizes and complexities. The architecture at each level mirrors the corresponding architecture used in the Animation Analyzer. The key difference is that the Pattern Recognizer does not use time-distributed wrapper, because it processes static pattern images rather than time sequences.

#### Decision Maker

The Decision Maker module consists of a concatenation layer followed by two fully connected dense layers. The concatenation layer merges the outputs from the Animation Analyzer and Pattern Recognizer into a single 1-D vector, which is then passed to the first dense layer. The output of this layer is subsequently fed into the second dense layer, where a Softmax activation function is used to compute the probabilities of behavioral categories.

The number of nodes in the first dense layer is determined by the higher complexity level between the Animation Analyzer and the Pattern Recognizer. This design allows the Decision Maker to adapt dynamically to the architectures of the other two modules. A batch normalization layer and a dropout layer (dropout rate = 0.5) are applied before the second dense layer to improve training stability and reduce overfitting.

### Detector training in *LabGym*

Adapted from *Detectron2*, an instance-segmentation package, the Detector is an integrated object tracking component of *LabGym*, efficient at object detection and preventing identity switches during social proximity. In the Generate Behavior Examples submodule of the Training Module, users may choose to use a trained Detector to detect animals and generate corresponding behavior examples for training a Categorizer. In the Analysis Module of *LabGym*, users may also choose to implement a trained Detector to assist the Categorizer in detecting objects and animals of interest during behavioral analysis. More details on the mechanisms of the Detector have been reported previously (Goss et al., 2024).

Like Categorizers, Detectors must be trained. The Detector used to track worms and generate the behavioral data reported in this study, *worm detector 9*, was trained using the Train Detectors submodule embedded within the Training Module of *LabGym*. To train, images were generated from multiple behavioral videos using the “Generate Image Examples” feature of the Train Detectors submodule, and individual worms from the multi-worm images were annotated and defined as a single class “worm” in *Roboflow* (Dwyer, B. et al., 2026). 433 images were annotated and augmented in *Roboflow* via horizontal/vertical flipping and brightness adjustment of ±10%. 1433 post-augmented images were then used for training the Detector with an inferencing framesize of 960 and 10,000 training iterations.

### Categorizer training in *LabGym*

One categorizer trained in *LabGym* was used to analyze the worm behavioral data reported in this study (*worm locomotion categorizer 22*). The Categorizer was trained using the Train Categorizers submodule embedded within the Training Module of *LabGym*. Training examples were augmented via five methods built into *LabGym*’s Train Categorizers submodule: random rotation, horizontal flipping, vertical flipping, random brightening, and random dimming. The Categorizer was trained on behavior examples generated in the “Non-interactive” mode, specified to identify behaviors of non-interacting individual worms. The Categorizer incorporated both the *Animation Analyzer* (complexity level 2) and *Pattern Recognizer* (complexity level 3) submodules. The training settings that resulted in the highest average macro categorization accuracy were retained: Animation Analyzer (LV2, 16 x 16 x 1) and Pattern Recognizer (LV3, 32 x 32 x 3). The performance metrics of each trained Categorizer, including categorization accuracy, precision, recall, and f1 score, were calculated on the Categorizer’s ground-truth testing data (sorted behavior examples) using the Test Categorizers submodule embedded within the Training Module. The descriptions and specific methods of calculating each type of performance metric have been previously reported in detail (Goss et al., 2024; Hu et al., 2023).

The hyperparameters used for training the Categorizer have been described previously (Hu et al., 2023); the key details are summarized below. Briefly, *LabGym* employs an automatic training pipeline that reduces the burden on users to manually set hyperparameters. Instead, users only need to focus on sorting behavior examples, while *LabGym* automatically determines appropriate training hyperparameters based on the amount of training data and the progress of training.

The loss function used to train the Categorizer is binary cross-entropy when the number of behavioral categories is two, and categorical cross-entropy when the number of behavioral categories is three or more. Stochastic gradient descent (SGD) with an initial learning rate of 1×10^−4^ is used as the optimizer during training. The learning rate is reduced by a factor of 0.2 if the validation loss stops decreasing (a decrease of < 0.001 is considered “no decrease”) for two training epochs. Training is terminated if the validation loss does not decrease for four consecutive epochs. The model is automatically updated whenever a training epoch achieves a new minimum validation loss.

The batch size used during training is automatically determined according to the size of the validation dataset: 8 when the number of validation samples is fewer than 5,000, 16 when it is between 5,000 and 50,000, and 32 when it exceeds 50,000. The dataset is split into training and validation subsets with a ratio of 0.8:0.2.

To improve training efficiency and enhance the generalizability of the trained models, we implemented nine data-augmentation methods that apply identical random transformations to both an animation and its paired pattern image. Importantly, during data augmentation, all transformations are applied only to the animal foreground and its contour; the absolute dimensions of the frames/images and the black background remain unchanged. These augmentation methods can be applied in combination, and users can choose which methods to use. When all augmentation methods are enabled, the dataset size can be expanded to approximately 200 times the original amount.

The data-augmentation methods are as follows:

1. **Random rotation.** Animations and their paired pattern images are rotated by a random angle within the ranges 10°-50°, 50°-90°, 90°-130°, 130°-170°, 30°-80°, or 100°-150° (the latter two ranges are used in combination with random brightness changes).
2. **Horizontal flipping.** Animations and their paired pattern images are flipped horizontally.
3. **Vertical flipping.** Animations and their paired pattern images are flipped vertically.
4. **Random brightening.** The brightness of animations is increased by a random value between 30 and 80.
5. **Random dimming.** The brightness of animations is decreased by a random value between 30 and 80.
6. **Random shearing.** Animations and their paired pattern images are sheared with a random factor in the ranges -0.21 to -0.15 or 0.15 to 0.21.
7. **Random rescaling.** Animations and their paired pattern images are rescaled in either width or height with a random ratio between 0.6 and 0.9.
8. **Random deletion.** One or two frames in the animations are randomly removed and replaced with black images of the same dimensions.

### Criteria for sorting the behavior examples

The Categorizer *worm locomotion categorizer 22* analyzed N2 worm locomotion, and outputs were used to compare *LabGym*’s accuracy with manual analysis. To train the Categorizer, we sorted “Non-interactive” mode behavior examples (12 frames in duration) into behavior categories. Behavior examples were extracted from 4 fps (frames per second) videos of D1 N2 worm locomotion and 4 fps videos of aging N2 worm locomotion using the *worm detector 9* in the Generate Behavior Examples submodule. The criteria and number of behavior examples used to train each behavior category are described as follows (1 example pair consists of one Animation and its corresponding Pattern Image):

*Forward crawling* (707 example pairs): The worm crawls in the forward direction, where sinusoidal body waves propagate from the tail toward the head. Typically slower and longer in duration than reversals.

*Reverse crawling* (788 example pairs): The worm crawls in the reverse direction, with typically deeper sinusoidal body waves than forward locomotion that propagate from the head toward the tail. Typically brief and often succeeded by an omega bend.

*Omega bend* (563 example pairs): The worm exhibits a deep, omega-shaped body bend where the head touches the tail, typically followed by a resumption of forward crawling in a new direction.

*Immobile* (400 example pairs): The worm completely ceases active movement/locomotion. *Twitching* (428 example pairs): The worm displays rapid, small-amplitude body sweeps usually localized in the head and/or tail.

Behavior examples included adult N2 worms that were 1-day-old, 3-day-old, 5-day-old, 7-day-old, 9-day-old, 11-day-old, 13-day-old, and 15-day-old. The quantity of behavior example pairs in the training dataset, broken down by age, is as follows:

D1 (1,478 example pairs), D3 (115 example pairs), D5 (141 example pairs), D7 (249 example pairs), D9 (230 example pairs), D11 (169 example pairs), D13 (242 example pairs), and D15 (262 example pairs).

### Categorizer Testing

The performance metrics of Categorizers are internally tested in *LabGym* via the Test Categorizers function unit, which uses the user-established ground-truth testing dataset to predict behavioral categories in a simulated environment (See Methods in Goss et al., 2024; Hu et al., 2023). The testing dataset consisted of the same behavioral categories as the training dataset, but the behavioral examples in each category were not included in the training dataset. The number of examples used to test each behavior category are described as follows:

*Forward crawling* (238 example pairs)

*Reverse crawling* (179 example pairs)

*Omega bend* (164 example pairs)

*Immobile* (125 example pairs)

*Twitching* (149 example pairs)

The following prediction equations were used to calculate performance metrics of Categorizers, which included precision (Equation 1), recall (Equation 2), f1 score (Equation 3), and overall accuracy (Equation 4).

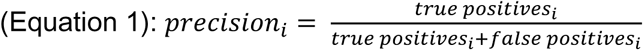

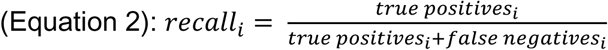

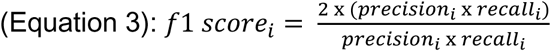

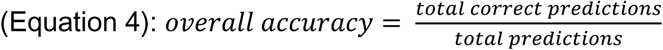

### Confusion matrix (Figure 2b)

The confusion matrix was generated using scikit-learn library in Python 3.10. The Categorizer *worm locomotion categorizer 22* was used to generate predicted behavior labels and the true labels were generated from the testing dataset described in the Categorizer Testing Methods section, which was also used to test the Categorizer.

### Ablation analysis (Figure 2c)

To perform the ablation analysis, the Categorizer was modified by either ablating the Pattern Recognizer (PR) or Animation Analyzer (AA) module. We then trained 3 types of Categorizers, Categorizer with AA only, Categorizer with PR only, and Categorizer with both, on the same training dataset using the same network setting (complexity level 2, input size 16). We then test their performance on the same testing dataset.

### *C. elegans* behavioral analysis

#### Analysis of D1 N2 locomotion using *LabGym* (Figure 3)

The Categorizer *worm locomotion categorizer 22* analyzed D1 N2 worm locomotion, and outputs were used to compare *LabGym*’s accuracy with manual analysis. For behavior classification, we set the Categorizer at an uncertainty threshold of 5% and a minimum behavior episode length of 3 frames, a setting that maximally optimized the accuracy of behavioral identification. These settings were optimized from a trial-and-error approach: different values for the uncertainty level and minimum behavior episode length were tested, and the resulting behavior raster plot and annotated video were manually evaluated for accuracy. The settings that yielded the relatively best detection and classification were chosen.

We analyzed the first 2-minute window of four behavioral videos of Day 1 N2 worms, proportionally downsized from 2056 x 2056 pixels to a frame size of 1028 x 1028 pixels, with six worms in each video. For each of the six target behaviors the Categorizer was trained to identify (behavioral categories described previously in Methods), quantitative measurements were outputted via *LabGym*’s Quantifier Module. The specific calculation methods of the quantitative measurements have been previously reported in detail (Hu et al., 2023; Goss et al., 2024). Duration was the parameter that was manually scored and used in the benchmark comparison between the output of *LabGym* and manual analysis, as it was one of the few parameters that could also be scored manually (Figure 3c). Raster plots of the behavior events in the 2-minute analysis window were generated to depict the results of *LabGym* analysis and manual scoring (Figure 3a). *LabGym* behavior plot generation was automatic and derived from an *all_events* Excel sheet that records the behavior event and corresponding event probability for each individual at each analyzed frame (every 0.25 seconds).

#### Manual analysis of D1 N2 locomotion (Figure 3)

The same 2-minute time window of four behavioral videos of Day 1 N2 worms analyzed by *LabGym* was also analyzed by eye to conduct a benchmark comparison. We acquired the duration of each behavioral event and the corresponding time of occurrence in the videos. To generate the raster plot summarizing behavior events scored from manual analysis, the manually-scored behavior events at each frame (every 0.25 seconds) were recorded) in an Excel sheet, formatted similarly to the *LabGym*-generated *all_events* Excel sheet. This Excel sheet was then used to generate a behavior plot via *LabGym*’s built-in Generate Behavior Plot submodule, embedded within the Analysis Module.

#### *WormLab* analysis of D1 N2 locomotion (Figure 3)

The same 2-minute time window of four behavioral videos of Day 1 N2 worms analyzed by *LabGym* was analyzed by *WormLab* (version 4.0.5) to conduct a benchmark comparison. For Detection and Tracking, Crawling Mode and back tracking were chosen, with a Max tracked hypotheses of 2. A Detection frequency of 10 frames was set, as well as 200 for Fitting iterations of the Worm Shape. Default settings were used for all other parameters.

To generate the raster plot summarizing behavior events scored from *WormLab* analysis, the *WormLab* behavior events at each frame (every 0.25 seconds) were recorded in an Excel sheet, formatted similarly to the *LabGym*-generated *all_events* Excel sheet. This Excel sheet was then used to generate a behavior plot via *LabGym*’s built-in Generate Behavior Plot submodule, embedded within the Analysis Module.

#### Quantification of analysis error during multi-worm collisions

To quantify analysis errors during multi-worm tracking by *LabGym*, we manually quantified the occurrences of Categorizer error during the collision events of multiple worms. Errors fell into two categories: 1. Detection loss: temporary loss of worm ID during interaction, with subsequent reidentification post-interaction, and 2. Behavioral misclassification: incorrect categorization of behavioral type during collision.

#### Analysis of locomotion of aging worms using *LabGym* (Figure 4)

The Categorizer *worm locomotion categorizer 22* was used to analyze behavioral changes of aging N2 worms from D1 to D15 adults (Figure 3). For behavior classification, we set the Categorizer at an uncertainty level of 5% and a minimum behavior episode length of 3 frames. We analyzed the full 4.5-minute duration of seven behavioral videos of D1, D3, D5, D7, D9, D11, D13, and D15 adult worms. Three videos per age were analyzed, and each video was proportionally downsized from a frame size of 2056 x 1716 pixels to 1028 x 858 pixels and contained 9-14 worms. For each of the six target behaviors the Categorizer is trained to identify (behavioral categories described previously in Methods), quantitative measurements were outputted via *LabGym*’s Quantifier Module. Forward crawling average speed, Forward crawling distance, Reverse crawling frequency/count, Omega bend frequency/count, Immobile duration, and Twitching duration were compared across ages (Figure 3b-g).

#### Trajectory Images

Images of worm trajectories, color-coded by individual, are automatically generated by *LabGym* (Figure 3a). To output images of worm trajectories color-coded by time instead of by individual animal (Figure S1), we utilized the Generate Behavior Examples submodule to generate a single behavior example pair (Animation + Pattern Image) on “Interactive Basic” mode (examples contain all detectable animals/objects in the behavioral video) for a video from each of the seven worm ages. To ensure the output of only one single behavior example pair that summarized the entire behavioral video, we first reduced the frame rate of each behavioral video from 4 fps to 1 fps via the Preprocessing Module, then inputted the total number of frames in the entire behavioral video when prompted to specify how many frames for each generated behavior example in the Generate Behavior Examples submodule. The Pattern Image from the behavior example pair for each video, now representative of the movement patterns of all worms over the entire duration of the behavioral video, was then used as the trajectory images (Figure S1).

### Statistical Analysis

All statistical analyses were performed using GraphPad Prism 10.0 software. Statistical methods, error bars, and p-values are indicated in the figure legends. Two-way ANOVA with Bonferroni multiple comparisons post hoc analysis and additional tests were applied to analyze data, specifics further described in the figure legends. For all statistical analyses: ns for p>0.05, * for p<0.05, ** for p<0.01, *** for p<0.001, **** for p<0.0001.

## ACKNOWLEDGEMENTS

We thank Shuhao Wan for valuable comments. Strains from the Caenorhabditis Genomics Center (CGC) were used in this study.

## STUDY FUNDING

This work was supported by grants from NIH (R01 NS128500 and R01 DC018167) and the LSI Innovation Partnership Fund to B.Y. and T32 DE007057 and T32 DC00011 to E.A.R.

## AUTHOR CONTRIBUTION

B.Y., Y.H., E.A.R., L.A.X., and X.Z.S.X. conceived the project. H.L., Z.L., and E.A.R. recorded behavioral videos. L.A.X. labeled datasets, trained the Categorizers and Detectors, and analyzed behavioral videos using *LabGym*. E.A.R. analyzed behavioral videos using *WormLab*. Y.H. designed and programmed *LabGym*. L.A.X., B.Y., Y.H., and E.A.R. wrote the manuscript with assistance from all other authors.

## COMPETING INTERESTS

The authors declare no competing interests.

## DATA AND CODE AVAILABILITY

An extended user guide with instructions on how to install and use *LabGym*, as well as the software’s open-source code, is publicly available on the *LabGym* GitHub page: https://github.com/umyelab/LabGym. The *LabGym* models reported in this study, as well as the behavioral datasets used to train them, are accessible on Zenodo (https://doi.org/10.5281/zenodo.19562601).

---

**Figure S1.**
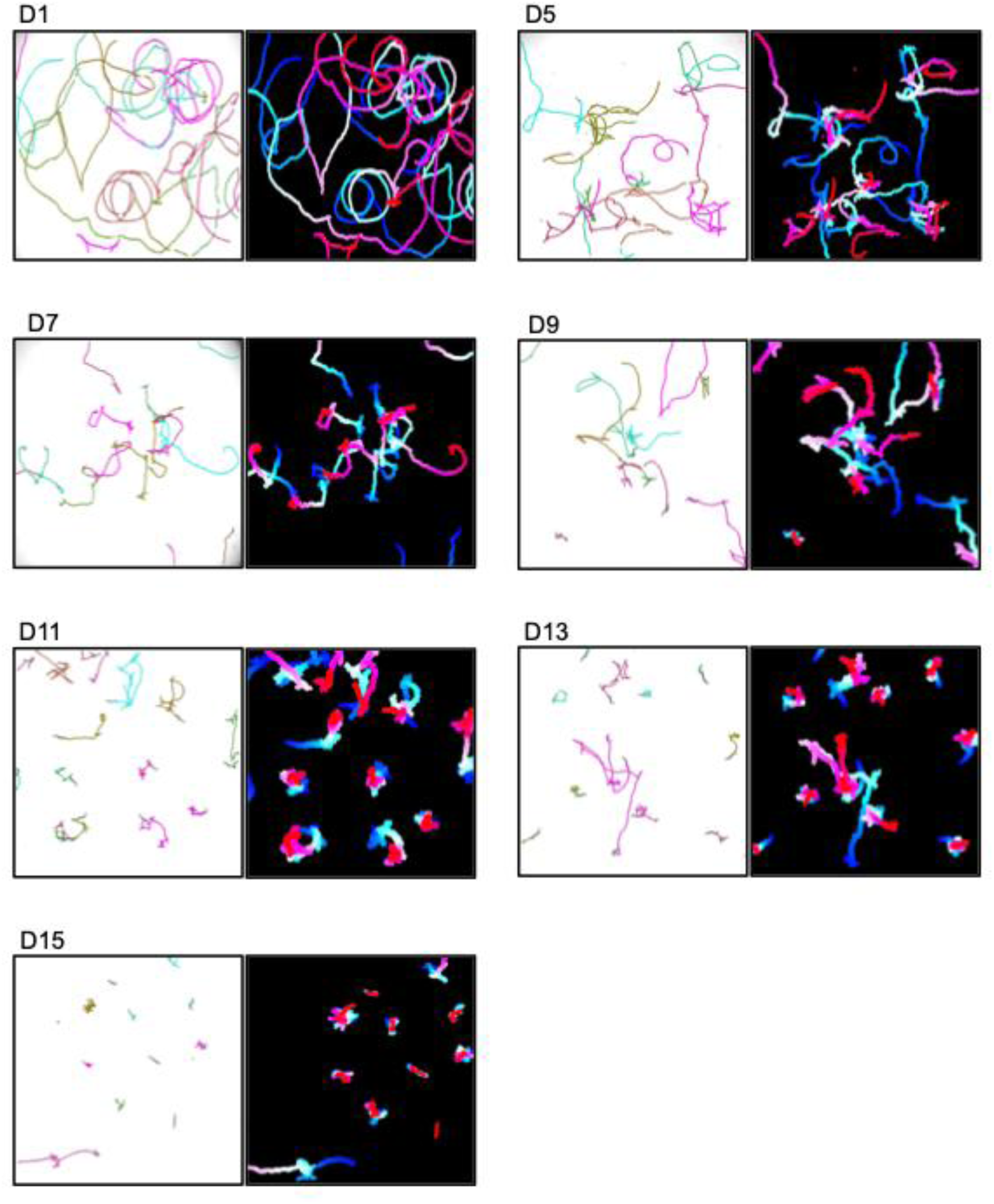
Locomotion trajectories of aging worms, color-coded by individual and by time. For each age, a pair of trajectory images was generated to depict individual worm movement paths, color-coded by individual (left image) or time (right image). For trajectory images color-coded by individual, different colors indicate different individuals. For trajectory images color-coded by time, color gradients indicate temporal progression of movement (earlier timepoints in cool tones, later timepoints in warm tones). These visualizations highlight age-dependent changes in the activity patterns and movement trajectories of worms across adult lifespan.

**Table S1.**
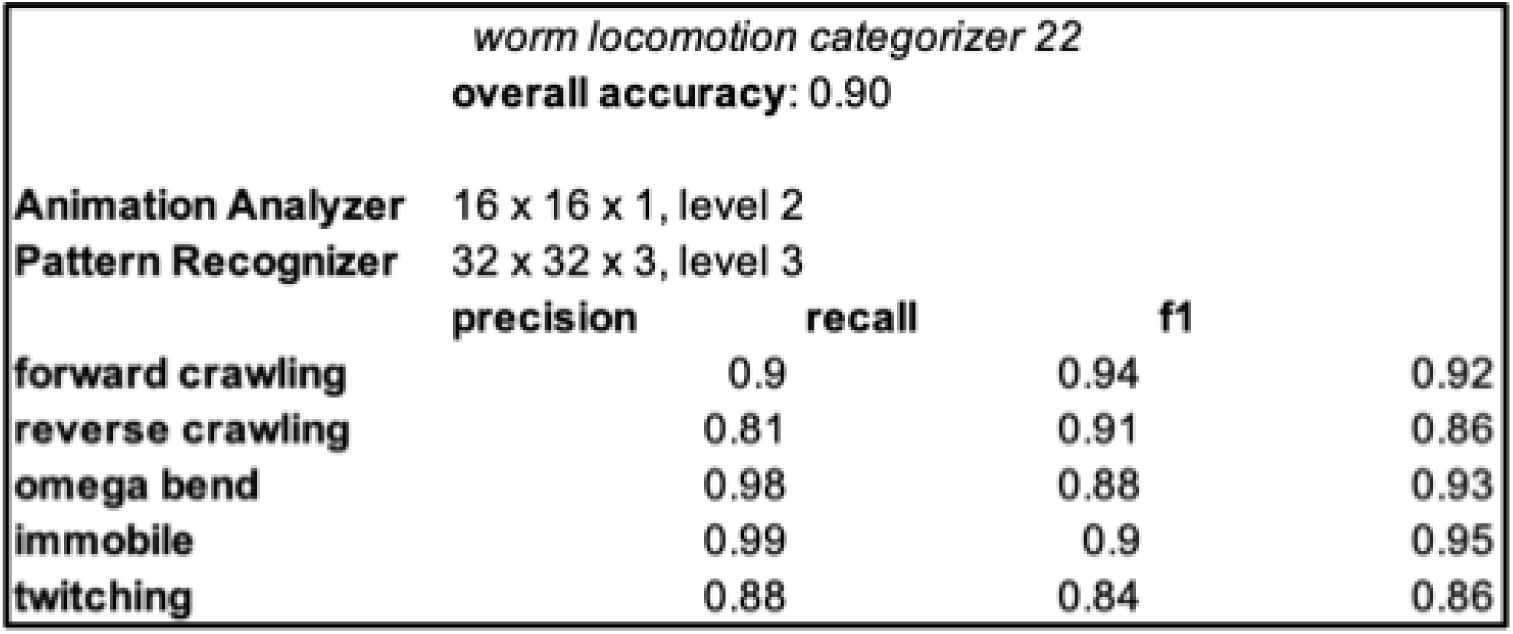
Summary Table of trained Categorizer metrics. Table summarizing the metrics of the trained Categorizer (overall accuracy, Animation Analyzer, Pattern Recognizer, and every behavior category’s precision, recall, and f1-score).

**Video S1. Representative behavior examples used for Categorizer training**

Representative behavior examples used to train the Categorizer. The sorting criteria and total number of example pairs for training each behavior category are detailed in Methods.

**Video S2. Annotated videos of N2 worm locomotion by *LabGym***

**Left:** Behavioral video of six Day 1 adult N2 worms with frame-by-frame annotation of computed behavior category and associated probability. 5x speed, 20 frames per second.

**Right:** Behavioral video of six Day 1 adult N2 worms with frame-by-frame annotation of computed behavior category and associated probability. 5x speed, 20 frames per second.

**Video S3. *LabGym* maintains detection accuracy during multi-worm collision**

A video example of a multi-worm collision event where *LabGym* maintains analysis accuracy before, during, and after the collision of two forward crawling worms without identity switching. Although *LabGym* is capable of differentiating individual worms when they physically contact each other in most scenarios, it must be noted that the Categorizer may miscategorize behaviors during and/or a few frames post-collision with an error rate of ∼30%, but can typically correct itself relatively quickly following entanglement. Given the Categorizer’s overall accuracy of ∼90%, such miscategorizations during collisions account for much of the remaining ∼10% inaccuracy. Retraining the Categorizer with more behavior examples involving multi-worm collisions is likely to improve its accuracy.

## Notes

https://doi.org/10.5281/zenodo.19562601

### Competing Interest Statement

The authors have declared no competing interest.

### Summary of Updates

The following is a list of major changes added to the revised manuscript. 1. We have included new image and behavior examples from independent trials across all adult age groups for retraining an all-encompassing Detector and subsequently a comprehensive Categorizer. 1433 image examples were used to train the new Detector, worm detector 9, and 2,886 behavior example pairs (nearly double the original) were used to train a new comprehensive Categorizer, worm locomotion categorizer 22. Each behavior type had a strict minimum of 400 examples. Quantitative assessments confirmed improved accuracy of the new Categorizer. 2. We added an ablation analysis (Figure 2c) to quantify the contributions of Pattern Recognizer and Animation Analyzer. 3. To account for plate-level variation, we expanded our external validation to include behavior videos from separate plates recorded on different days for all experimental groups. 4. We have expanded the Introduction and Discussion to contextualize LabGym within the current open-source landscape (e.g. Tierpsy Tracker, WormPose, and DeepPoseKit). Specifically, we highlight the skeleton-free, holistic assessment of LabGym as a user-friendly solution to common worm analysis bottlenecks that bypass the need for explicit head/tail assignment. 5. We have included a Supplementary Quick-Start Guide and detailed hardware specifications to ensure ease of reproducibility for our methodologies. We also confirmed the public availability of our models and training datasets on Zenodo.

