## Supplementary material for "Automated analysis of *C. elegans* behavior by *LabGym*: an open-source, AI-powered platform": Supplemental File1_Quick-Start Guide.pdf

This quick-start guide provides a concise overview of the workflow used in this study to optimize *LabGym* for analyzing and quantifying *C. elegans* locomotion. The trained Categorizers and Detectors used in this work are publicly available and can be downloaded from Zenodo (https://doi.org/10.5281/zenodo.19562601)

For more detailed instructions, tutorials, and tips on troubleshooting, please refer to the published *LabGym* tutorials and user guides on the *LabGym* GitHub and website:

- GitHub: https://github.com/umyelab/LabGym
- Website: https://labgym.org

### Workflow Overview

1. Acquire Behavioral Videos
   Record videos of *C. elegans* performing behaviors of interest.
   - Use a frame rate and resolution sufficient to capture fine locomotor details and body parts, if applicable. This may vary depending on the behaviors/animals of interest.
     - An original frame rate of 8 fps and resolution of 2056 x 2056 pixels was used to capture the original worm behavioral videos analyzed in Figure 3 of this paper (Video S2 and Figure 3).

2. Preprocess Videos in *LabGym*
   Use the Preprocessing Module to optimize video properties for generating Categorizer training data:
   - Some common video properties to adjust include frame rate, resolution, contrast, and brightness.
     - The worm behavioral videos analyzed in Figure 3 of this paper were adjusted to a frame rate of 4 fps, downsized to a resolution of 1028 x 1028 pixels, with both the contrast and brightness increased 1.4 fold (Video S2).
   - If videos include extraneous features, one can also crop video frames to region of interest and trim video duration to only include time windows of relevance/interest.
     - The behavioral videos analyzed in Figure 3 of this paper were trimmed down from an original duration of 2 minutes and 40 seconds to 2 minutes.

3. Train Detectors
   - Generate Image Examples to extract representative frames from behavioral videos containing animals of interest. Annotate images by identifying objects of interest (recommended annotation tools include Roboflow or LabGym's built-in annotation module, EZannot) to train Detectors for recognizing the worms or animals/objects of interest.
     - The Detector reported in this paper, *worm detector 9*, was trained on 1,433 image examples featuring worms extracted from behavioral videos captured under different magnifications and settings. To train the Detector, an inferencing framesize of 960 and training iterations of 10,000 were implemented.

- Evaluate Detector accuracy using the Test Detector module. To build a testing dataset for accuracy assessment, select additional representative images from diverse settings (around ¼ the quantity of images in the training dataset) that are ideally images not used to train the Detector.
    - *worm detector 9* was trained on 1,433 image examples and tested on 368 image examples that were not used in training.

4. <u>Train Categorizers</u>
    - Generate Behavior Examples to extract pairs of animations and pattern images illustrating behaviors of interest (such as forward crawling, reversals, omega bending, etc. in worms). Then, sort behavior pair examples into different behavior categories to train Categorizers to recognize and quantify such behavioral types.
        - The Categorizer reported in this paper, *worm locomotion categorizer 22*, was trained on 5 behavioral categories (forward crawling, reverse crawling, omega bend, immobile, and twitching) with an accuracy of 0.9 (Figure 2a).

    - Evaluate Categorizer accuracy using the Test Categorizer module. To build a testing dataset for accuracy assessment, sort additional behavioral pair examples into behavioral categories identical to those in the Categorizer (around ¼ the quantity of examples in the training dataset) that are ideally behavior examples not used to train the Categorizer.
        - *worm locomotion categorizer 22* was trained on, across all behavioral categories, 2887 total example pairs and tested on 855 example pairs that were not used in training (Figure 2a).

5. <u>Analyze Behavioral Videos</u>
   Using the Analysis Module, apply the trained Detectors and Categorizers to automatically identify and quantify behaviors of interest in acquired behavioral videos *LabGym* outputs include:
    - Annotated videos with labeled behaviors (Video S2)
    - Quantitative summaries of each behavior (count, latency, duration, velocity, etc.) (Figures 1e and 3c)
    - Time-aligned raster plots of behavioral events (Figure 3a)
